# Evolution of explorative and exploitative search strategies in collective foraging

**DOI:** 10.1101/2023.01.03.522515

**Authors:** Ketika Garg, Paul E. Smaldino, Christopher T. Kello

## Abstract

Evolutionary theories of foraging hypothesize that foraging strategies evolve to maximize search efficiency. Many studies have investigated the central trade-off between explore-exploit and how individual foragers manage it under various conditions. For foragers in groups, this trade-off can be affected by the social environment, influencing the evolution of individual search strategies. Previous work has shown that when learning socially, explorative search strategies can optimize group search efficiency. However, social learning can cause discrepancies in strategies that benefit the group vs. an individual. We model the evolution of explorative and exploitative strategies using Lévy exponents under different levels of social learning and investigate their effect on individual and group search efficiencies. We show that reliance on social learning can lead to the evolution of mixed groups that are not optimally efficient. Exploiters can have a selective advantage in scrounging findings by explorers, but too many exploiters can diminish group efficiencies. However, greater opportunities for social learning can increase the benefits of explorative strategies. Finally, we show that area-restricted search can help individuals balance exploration and exploitation, and make groups more efficient. Our results demonstrate how exploration and exploitation must be balanced at both individual and collective levels for efficient search.

## 1 Introduction

Collective search, where independent individuals share information with each other to search for resources in a physical landscape, or for solutions in a problem-space, is common across biological and social contexts. Collective searching can decrease search costs, efficiently exploit resources, make search reliable, and enhance sensing of environmental information [Clark and Mangel, 1986, Krause et al., 2002, Berdahl et al., 2013, Harpaz and Schneidman, 2020, Roeleke et al., 2022, Aplin et al., 2014, Lihoreau et al., 2016]. It may therefore benefit individual searchers and their individual success to be part of a group [Giraldeau and Caraco, 2018, Pitcher et al., 1982]. However, competition for limited resources can make information-sharing costly [Ranta et al., 1993] and result in discrepancies between search behavior that is adaptive for the individual versus the group [Leonard and Levin, 2022]. In this study, we investigate the evolution of individual search strategies under the constraints and affordances of collective foraging, how they affect collective search, and what factors may facilitate the evolution of strategies that are beneficial for individuals as well as groups.

To efficiently search, individual foragers must balance exploration for new resources with exploitation of known resources [Kamil et al., 1987, Bartumeus et al., 2014, 2016, Hills et al., 2015, Garg and Kello, 2021]. Previous works have shown that efficient searchers need to modify this balance based on their environments (such as resource density and availability [Charnov and others, 1976, Krebs et al., 1978, Bartumeus et al., 2014, 2016]). This balance may further change when the search occurs in a group where individuals can interact and affect each other’s strategies. For example, foragers may aggregate to collectively exploit a found patch of resources, but too much aggregation may increase competition between foragers [Krause et al., 2002] and heighten the need to independently explore for new patches. Previous studies from animal foraging have shown that foraging behaviors in individuals, in terms of exploration tendencies or boldness, can be affected by social competition [Bergmüller and Taborsky, 2010, Webster and Ward, 2011]. The degree to which foragers influence each others’ search strategies can depend upon how they interact, for example, through social learning.

Social learning, by which foragers can observe and acquire information about resources using social cues, can influence how beneficial an explorative (or exploitative) strategy is. The advantages of a strategy under social learning can further depend upon the properties of the physical and social environment, such as resource density and group size. Herein, we develop a minimalist evolutionary model of collective foraging that combines independent foraging with social learning to investigate how individual-level search strategies can be shaped in response to constraints imposed by social learning and the foraging environment. We also investigate the effects of evolved individual strategies on group-level search performance and which strategies may maximize the benefits of collective foraging. We aim to simulate basic interactions between individual search and social learning that can hold general implications for bio-social systems such as animal groups, human teams [Toyokawa et al., 2014, Giannoccaro et al., 2020, Alexander et al., 2015], and multi-robot swarms [Lerman and Galstyan, 2002, Winfield, 2009] where individual agents in an information-sharing group can differ in their explore-exploit tendencies for independent search to find resources or solutions.

Social learning can influence individual foraging decisions to explore or exploit[Strandburg-Peshkin et al., 2017, Spiegel and Crofoot, 2016, Sokolowski et al., 1997, Greene et al., 2016]. For example, if foragers can detect when and where others find resources, foragers may forego costly, independent exploration and instead prefer to engage in social learning and head towards locations where others have found resources in the hopes of finding more nearby. However, socially driven search strategies may also increase competition for resources and cause foragers to adopt more explorative strategies that help them distribute themselves in ways that counteract a tendency to over-aggregate and over-exploit found resource locations [Beauchamp, 2005, Gillespie and Chapman, 2001, Di Bitetti and Janson, 2001]. The additional costs of collective foraging may necessitate foragers to inform their search decisions by the current state of the environment, for example, through area-restricted search [Pacheco-Cobos et al., 2019, Hills et al., 2013, Kerster et al., 2016, Hecker and Moses, 2015]. The effects of social learning may also interact with other search conditions such as group size and group composition to modulate the level of social information available. An increase in group size can amplify social information (for example, in bees [Detrain and Deneubourg, 2008]), especially in rich environments, and decrease overall exploration.

However, reliance on social learning may lead individuals to adopt search strategies that are not adaptive at the group level. In a previous study [Garg et al., 2022], we showed that foraging groups could maximize their search efficiency when individual foragers independently explore the environment while selectively joining other foragers in their discoveries. However, at the individual level, such explorative strategies may not always be adaptive, especially if exploitative agents can decrease their search costs in the presence of other explorative foragers. Previous models on group foraging have shown that certain foraging strategies like producer or scrounger can be frequency-dependent, i.e., a strategy’s pay-off depends upon the frequency at which it is adopted [Barnard and Sibly, 1981, Vickery et al., 1991, Afshar and Giraldeau, 2014]. Theoretical and empirical works have shown that such frequency-dependent dynamics can also prevent populations from evolving to group-optimal equilibria [Rogers, 1988, Smith and Price, 1973, Turner and Chao, 1999, Svensson and Connallon, 2019, Nowak and Sigmund, 2004].

In the present study, we investigate how reliance on social learning can shape the evolution of individual search strategies along the exploration-exploitation continuum. We also test the effects of these evolved strategies on collective search efficiency, i.e., group fitness. In addition, we study how individual-level search strategies can evolve to be beneficial for the group and the individual. In the previous model ([Garg et al., 2022]), all agents in a group practiced the same search strategy and we calculated which strategy maximized collective efficiency. Here, agents could vary in their degree of exploration versus exploitation by means of a parameter *μ* that governs the probability of relatively short versus long movements through the resource landscape. Using an evolutionary algorithm, *μ* parameter values were selected based on their effects on individual efficiency and in the context of different levels of social learning. Social learning was governed by a group-level parameter, *α* that determined how likely agents were to use social information to find resources. The model does not consider the extent to which social learning (*α*) evolves but rather considers the downstream evolutionary consequences of a population that relies more or less on social information. We would also like to note that our model studies selection for efficient search at the individual level, instead of the group. Our results show that reliance on social learning can lead to frequency-dependent dynamics between exploration and exploitation. Our results also show that group performance declines when social learning promotes the evolution of exploitative strategies. Finally, we show how informed search strategies that effectively balance exploration and exploitation can counteract the over-reliance on social learning by improving the payoffs of explorative search.

## 2 Model Overview

We developed an evolutionary model of collective foraging under different conditions of resource density, group size, and social learning that could constrain the evolution of individual search strategies. In the model, agents are conspecifics foraging for resources in patchy environments and their fitness is based on search efficiency. Agents can search for resources independently or they can learn about resource locations discovered by other foragers. Each simulation of the model was run until 30% of the available resources were consumed, at which point all resources were cleared and refreshed, and all agents were replaced by a new generation. The new generation of agents was copied from the previous generation in proportion to the fitness of each previous agent. The new generation thereby inherited their parents’ search strategies, such that the persistence of a search strategy depended on its efficiency given the search conditions.

We characterize exploration by how extensive (or long-range) an agent’s search is, which is correlated with their probability of finding new resources and the time they spend near an already discovered resource. To simulate different levels of exploration, we adapted the Lévy walk model [Viswanathan et al., 1999b] which can generate a range of random search strategies along the exploration-exploitation continuum. The model uses a power-law parameter *μ* that modulates the probability of relatively long versus short search movements and thereby simulates observed features of explorative and exploitative search behaviors [Mehlhorn et al., 2015]. For example, agents with *μ* → 1 employ a relatively large proportion of long, straight movement steps that help to cover new ground and find new patches at a faster rate compared with shorter steps that are more likely to double back on themselves. Long movements can reduce excessive overlap between agents but they may also cause agents to exit a patch before fully exploiting its resources. The Lévy walk model was chosen in part because a single parameter controls the variation between highly exploitative and highly explorative search strategies.

We also test a model of informed search, where foragers can adaptively switch between exploration and exploitation based on information about the distribution of resources in their environments. To do so, we modified the model to include area-restricted search (ARS), which is a simple heuristic that triggers slower and exploitative movements after encountering resources to search for more nearby before reverting to more wide-ranging explorative movements. ARS may increase the individual payoffs of explorative strategies and thereby counterbalance any social learning bias towards exploitative scrounging behaviors.

## 3 Model Details

Similar to the model of collective foraging from Garg et al. (2022), the search space is a *L* x *L* grid with periodic boundaries. Resources in each simulation were clumped into 20 randomly distributed patches under all conditions, and the total number of resources was varied to create either sparse or dense resource patches, *N*_*R*_ = 1000 or *N*_*R*_ = 10000 (see Supplementary Methods for more details and Fig. S1 for an example of the resource environment). Previous studies have shown that in environments with dense patches (where resources are easily available and present in higher numbers), foragers should prolong their time in a patch, i.e., be more exploitative. However, this effect of the resource environment may interact with the constraints posed by the social environment.

Resources in the model are destructive, i.e., they are removed from the environment after being found during each simulation. We would like to note that while resources in our model are destructive, they exist in clumped patches which can be revisited until exhausted. Thus, the patches simulate a form of non-destructive search [Viswanathan et al., 1999a] where there are usually resources available in the vicinity of the one found [Wosniack et al., 2017]. To ensure that such revisitable patch structures remain mostly intact throughout the course of the simulation, we end each generation when 30% of resources have been exhausted.

We tested the model under two different numbers of agents, *N*_*A*_ = 10 or *N*_*A*_ = 50. Group size (*N*_*A*_) varied the potential amount of social information available, and consequently the level of social interaction and competition among agents. Each agent foraged based on the following rules: On each time step, each agent consumes a resource unit if one existed within a radius, *r* = *d*_*min*_, or in other words, if a resource is present at their current grid location. Otherwise, the agent moves in search of additional resources. Heading and distance (*d*) of each move are chosen based on either the agent’s individual search strategy or the location of a detected social cue, i.e. where another agent found one or resources on the current time step. Each agent can detect all locations where resources are found by others on each time step. Agents execute each move of distance, *d*, in a series of steps of fixed size (*d*_*min*_ = 10^−3^), i.e., they have a constant speed and thus, longer steps take more time to execute and incur larger implicit costs.

To independently search, agents choose a random heading and move a distance sampled from the following probability distribution,

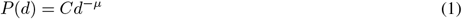

where, *d*_*min*_ ≤ *d* ≤ *L, d*_*min*_ = 10^−3^ is the minimum distance that an agent could move, *L* is normalized to the width of the grid, and *μ* is the power-law exponent, 1 *< μ* ≤ 3. *C* is a normalization constant such that

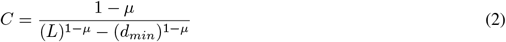

The Lévy exponent *μ* modulates the individual search strategy as a continuum between shorter, more exploitative movements and longer, more explorative movements. There were six different alleles i.e. possible values for *μ*: [1.1, 1.5, 2.0, 2.5, 3.0, 3.5] and each agent *i* is characterized by one of the exponent values. *μ* → 1.1 represents an explorative strategy with longer steps, *μ* → 3.5 results in an exploitative strategy with shorter steps and frequent turns, and *μ* ≈ 2 balances the probability of long versus short steps. Later, we consider ARS as an informed individual search strategy that can be added to parameterized Lévy walks.

An agent, *A*_*i*_, continues detecting social cues and resources at its current location at each time step while executing an independent search move (Lévy walk of total distance *d*). If there is no cue or resource available, it continues moving step by step (*d*_*min*_) in the direction of its previous heading until it finishes walking the distance, *d*. If the agent encounters a resource at its location, it consumes it and truncates its walk. If no social cue or resource is detected after executing the total distance, *d*, it draws a new heading direction and distance based on its Lévy exponent.

However, if any time step, it detects a social cue at a given location where another agent *A*_*j*_ has found a resource, then it truncates its current walk to pursue the social cue with the following probability:

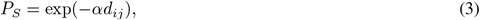

where *d*_*ij*_ is the distance between agents *A*_*i*_ and *A*_*j*_, and *α* is the social selectivity parameter that determines how selective an agent is in pursuing social information as a function of distance [Bhattacharya and Vicsek, 2014]. The agent continues detecting social cues at each time step while pursuing social cues or independently searching. If it detects a new social cue while following another cue, it pursues the new cue only if its distance is less than that of the currently pursued cue. If more than one social cue is detected on a given time step, the agent pursues the closest cue (with ties chosen at random).

We simulated the model for three levels of social selectivity (*α* = 10^−5^, 10^−2^, 10^0^) that correspond with high, intermediate, and no (or negligible) social learning, respectively. With high social learning, agents are more likely to pursue social cues irrespective of distance. With intermediate social learning, agents are more selective in pursuing social information and are less likely to pursue distant cues that incur greater movement costs. With no social learning, agents are very unlikely to follow social cues and instead forage using only their individual search strategy.

Agents consume resources at locations they encounter. If multiple resources are present at a given location, agents consume one unit of resource per time step, in the order of their arrival at the location. Thus, fewer or no resources are available for agents arriving relatively late to a given resource location. Agents truncate their movements (towards a social cue or random location) if they encounter a resource. After consuming all the resources at a location, they initiate a new move either towards a social cue or to independently search along a random distance and direction.

### Area-restricted search

Individual search strategies based on the above model are uninformed because steps are stochastic and unaffected by information that could be gained while foraging. We added ARS as an informed component of individual search strategies that is triggered when resources are found (similar to [Reynolds, 2009, Ross and Winterhalder, 2018, Hecker and Moses, 2015]). Specifically, when an ARS agent moves to a location with one or more resources, it searches the vicinity before moving to its next location, where vicinity was defined as all neighboring locations within a radius of two grid cells, *r*_*v*_ = 2*d*_*min*_. This radius is a proxy for intensive, local search upon encountering a resource that has been observed in various natural foraging conditions. ARS agents move to any of the neighboring locations where resources are found to consume them. ARS is potentially more efficient when resources are clustered and hence more likely to be near each other.

### Genetic Algorithm

Each simulation began with a group of agents with uniformly distributed values of *μ* and thus, groups represented the six alleles in equal proportions. The other three parameters (*N*_*R*_, *N*_*A*_, *α*) were held constant for each given simulation and varied systematically across simulations. The evolutionary algorithm selects agents based on their search efficiencies *η*,that represents the ratio of the benefit accrued and the total cost expended. We computed *η* as the total amount of resources consumed (benefit) per total distance moved (cost). Each round of selection occurs after 30% of resources are consumed, and efficiencies are normalized to assign each allele with a probability of replication proportional to efficiency.

Each new population inherits the Lévy walk exponent, *μ* from the selected parents. In addition, we added a mutation rate of 0.05 probability of randomizing *μ* to one of the six alleles for each agent on each selection round. Resources, efficiencies, and agent locations are reset after each selection round and the simulation continued anew. Our genetic algorithm selects individual strategies based on individual fitness, and not based on mean group fitness. We can expect the mean group fitness to increase over time with the intuitive hill-climbing dynamics if the fitness values of search strategies are independent of their frequencies. However, if the success of a strategy is dependent upon its relative frequency in the population, there is no guarantee that the overall mean fitness of the population will increase with time [Turner and Chao, 1999, Svensson and Connallon, 2019, Nowak and Sigmund, 2004].

### Evolutionary analyses

The results presented here are from 40 simulations, and each simulation was run for 3000 generations. We measured the evolved values of *μ* and group search efficiencies for each parameter combination and for both models (non-ARS and ARS). The results presented below show both the mean evolved *μ* in populations and their distribution across populations, mean search efficiencies (*η*), and the changes in *μ* and *η* over generations. Note that our model results in stochastic evolutionary dynamics due to variability in population sizes, resource environments, stochastic search decisions, spatial interactions, and mutations. Such stochasticity prevents the groups from evolving to fully stable equilibria [Imhof et al., 2005].

To corroborate our findings, we also performed invasion analyses with the model to test the likelihood of a strategy (*μ*_*mutant*_) invading a population of another strategy (*μ*_*resident*_) based on their relative payoffs (see Supplementary Methods for more details). The likelihood can be shown by calculating an invasion index (*i*) for a given set of resident and mutant strategies,

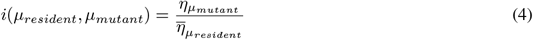

If a strategy is stable, then another strategy can not outperform it, and *i <* 1. If a strategy is unstable, then a mutant with a more efficient strategy can invade, and *i >* 1. If two strategies *μ*_*i*_ and *μ*_*j*_ can invade each other, the groups may evolve to stable, mixed equilibria with strategies *μ*_*i*_ and *μ*_*j*_ coexisting. The results from invasion analyses were not affected by parameters that might affect evolutionary simulations like the number of generations, mutation rate, and selection rules.

## 4 Results

### 4.1 Reliance on social learning leads to frequency-dependent payoffs and mixed groups of explorers and exploiters

Previous studies have shown that with little or no social learning (i.e., solitary foraging), individual search strategies that balance explorative and exploitative movements with the Lévy exponent, *μ* ≈ 2 are optimal for individual and group-level search efficiencies [Viswanathan et al., 2008, Garg et al., 2022, Bartumeus et al., 2016]. We similarly found that without social information, the genetic algorithm increased search efficiency by selecting individual search strategies with *μ* ≈ 2 (Figs. 1, (right column), 4 (top-left) and Fig. S2(right)). In line with existing literature [Wosniack et al., 2017, Bartumeus et al., 2014], we found that resource density affected the optimal balance between exploration and exploitation. Sparse patches necessitated more exploration of the search space and thus, tilted the balance more towards exploration relative to denser patches (see Fig. 1 (right-most column)).

**Figure 1:**
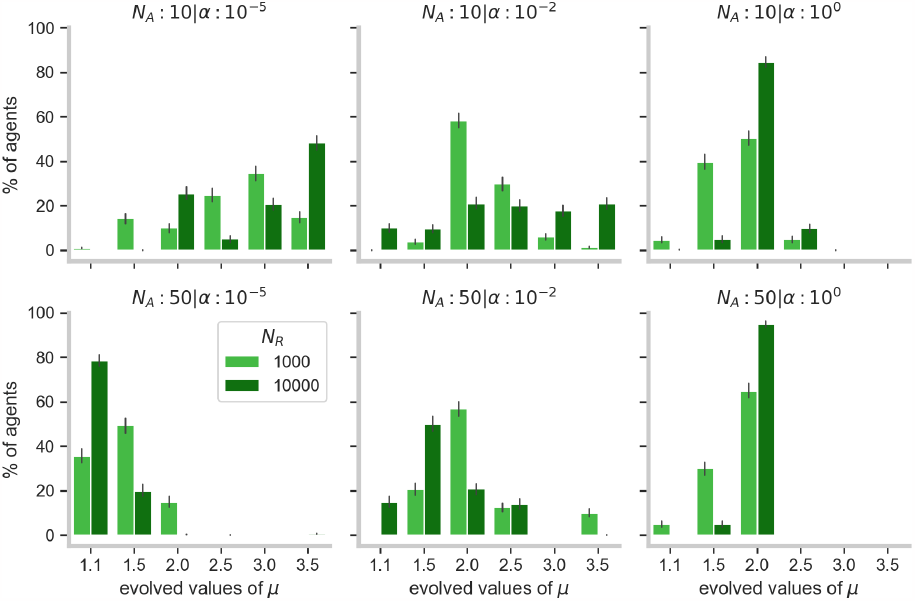
Distribution of the evolved values of Lévy exponents (*μ*) for different levels of resource density (*N*_*R*_), group size (*N*_*A*_) and social learning (*α*). *α* values of 10^−5^, 10^−2^, 10^0^ correspond to high, intermediate, and no levels of social learning. *μ* → 1 corresponds to an explorative search strategy, while *μ* → 3.5 corresponds to exploitative search. These data represent group compositions over the last 10 generations out of a total of 3000. Error bars show 95% confidence intervals.

Social learning can further modify the optimal balance between explore-exploit in individuals. Our previous model [Garg et al., 2022] showed that, at the group-level, social learning increased the efficiency of explorative search. The model showed that explorative search not only increases the rate of discovering new resources but also prevents excessive crowding at patches that can be a side-effect of too much reliance on social learning. However, in the current model, we found that when agents could use social information (*α <* 10^0^) to find resources instead of exploring independently, it was beneficial for them to adopt strategies that were more exploitative than explorative. In other words, although the explorative search is beneficial to a group that shares information, it does not confer a selective advantage to agents when selection operates at the individual-level. Instead, we found that agents evolved values of *μ* that fluctuated between explorative and exploitative strategies over generations, with an exploitative bias (see Fig. 1 (top-left)).

The fluctuation in distributions of individual search strategies with social learning can be explained by cyclical frequency-dependent dynamics. In the presence of a few explorative searchers who are quick to find resources, exploitative strategies become advantageous because they can scrounge off the explorers. But as exploitative strategies are increasingly selected, the proportion of explorative strategies drops and explorers become over-exploited. Without many explorers left to discover new resources, exploitative search becomes less efficient and the advantage swings back to explorative strategies, and so on. This cyclical dynamic is similar to negative frequency-dependent selection [Mottley and Giraldeau, 2000] that can lead to a mixed evolutionary stable strategy between explorers and exploiters: exploiters are more efficient when their frequency is low in the population and as a result, neither explorers nor exploiters can completely invade a population. We can also see this pattern in the invasion analyses (Fig. S4 (top-row)) where exploitative mutants could invade explorative populations, but exploitative populations could, in turn, be invaded by explorers with *μ* = 1.5.

The level of exploration/exploitation that individuals evolve to in a group, thus, depends upon the balance of payoffs from spending time searching for new resource patches vs. from prolonging time within patches. Social learning decreases the benefits of independently exploring and instead benefits exploitative strategies that can effectively scrounge others’ discoveries (more so with denser patches). But this benefit only holds until there are enough explorers to find new patches, at which point, it shifts back in favor of more exploration. We next ask how the payoffs from explore-exploit alter in large groups, where the opportunities for social learning increase and the costs of overall exploitation can exceed its benefits.

### 4.2 Greater opportunities for social learning in large groups increase the benefits of explorative strategies

Large groups of social learners can amplify the amount of social information produced and the opportunities to socially learn. Such widespread social learning can increase the group’s overall exploitation levels, which, in turn, can suppress the discovery of new resources and increase competition for the ones already found. We found that in large groups (*N*_*A*_ = 50), explorative search strategies were more likely to be selected than exploitative ones (Fig. 2a (right), Fig. 1 (bottom panel)).

**Figure 2:**
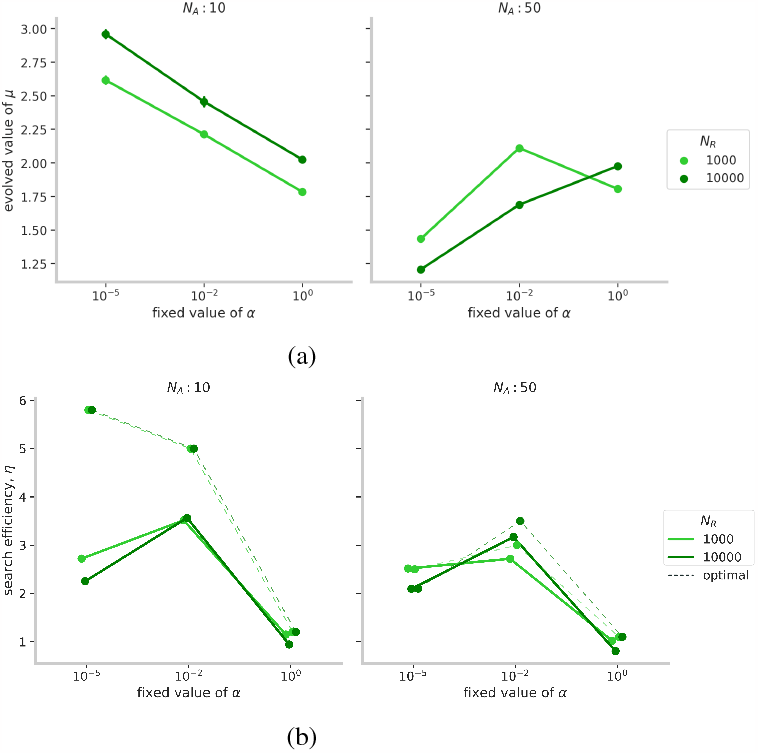
***(a)*** Mean estimates of the evolved values of Lévy exponents (*μ*) for different levels of resource density (*N*_*R*_), group size (*N*_*A*_) and social learning (*α*). ***(b)*** Corresponding mean estimates of the group search efficiencies (*η*) of the evolved groups. Dashed lines show the maximum group efficiency value obtained in Garg et al. (2022) for given *α* and *N*_*A*_. These values were similar for the two resource densities (*N*_*R*_). The averages were taken over the last 10 generations out of a total of 3000, for every parameter combination. Error bars indicate 95% confidence intervals.

The presence of more agents in a group led to more persistent social cues and caused the group to excessively aggregate at patches. Such overcrowding resulted in a quick depletion of patches while raising the level of competition between agents. As a result, exploitative strategies could not scrounge effectively and their payoffs decreased. Conversely, it was beneficial to adopt explorative strategies (*μ* → 1) that could not only quickly search the area extensively for new patches but they could also exit patches before their resources diminished. With low social selectivity (*α* = 10^−5^), a high proportion of explorative agents quickly covered the search space and coalesced at newly discovered patches before quickly breaking apart after exploiting its resources. On the other hand, exploitative agents, who could not find new resources independently, would not also be able to scrounge effectively because patches would be gone by the time of their arrival. High levels of competition for limited resources ensured that resources would not remain in a patch for too long after its discovery. Therefore, explorative strategies (*μ* ≈ 1.1) were less likely to be invaded by exploitative strategies (*μ* ≥ 2.5) under high levels of social learning (Figs. 1, S5).

Furthermore, when the competition was highest in the conditions with dense patches (*N*_*R*_ = 10, 000) and low social selectivity (*α* = 10^−5^), explorative strategies’ benefits increased and groups evolved to be quite explorative (Fig. 1). This effect is in contrast to the effect for small groups (see previous section), where dense patches meant sufficient resources for all agents at the patch and lessened the need for independent exploration.

Our results so far show that individual payoffs from how explorative (or exploitative) a strategy is vary based on the level of social learning. Individual foraging (i.e., no social learning) favors strategies that balance explore-exploit. Reliance on social learning makes groups overall more exploitative, but in large groups with greater opportunities to scrounge, explorative strategies are favorable to counterbalance the high levels of overall exploitation. We next test the effect of these evolved strategies on group-level efficiencies.

### 4.3 Selection for exploitative strategies due to social learning can make groups less efficient

Although our genetic algorithm was set up to select agents with high search efficiencies, we found that in many cases, group-level search efficiency actually decreased over generations (Fig. S2(left)). If many agents in a group evolved to be exploitative, group-level exploration dipped and groups performed substantially below the optimal levels as derived from the previously published model (Fig. 2b (left)). Conversely, when groups evolved to be explorative (in large groups, *N*_*A*_ = 50), groups maintained search for new resources and efficiencies did not diminish. We found that higher proportions of explorers corresponded with near-optimal search efficiencies (Fig. 2b (right)).

In small groups (*N*_*A*_ = 10), when agents used social learning, the presence of explorative agents decreased search costs faced by exploitative agents and made them more efficient than their explorative counterparts. This advantage allowed exploitative agents to invade the population, but the absence of explorers decreased their efficiencies and resulted in less efficient groups. We found that higher proportions of exploitative searchers at any time corresponded with lower efficiencies. For example, in Figs. 3a and 4 (top row), the mean *μ* of the group (shown in blue) decreased and increased periodically, and the decreases in *μ* coincided with elevated search efficiencies (shown in green). An increase in explorers made exploiters more efficient searchers and resulted in more efficient groups. Exploiters could then replace explorative agents but at the expense of mean efficiencies. This effect caused dips in mean efficiency to coincide with increases in mean values of *μ*. As a result of high levels of exploitation, evolved groups were substantially less efficient than the optimal search efficiencies predicted in Garg et al. (2022) (shown with dashed lines in Fig. 2b). This decrease in group search efficiency was more pronounced with rich resource patches (*N*_*R*_ = 10, 000), where exploitative strategies that prolonged the time spent within a patch were more advantageous than the strategies that left too early (Fig. 2a).

**Figure 3:**
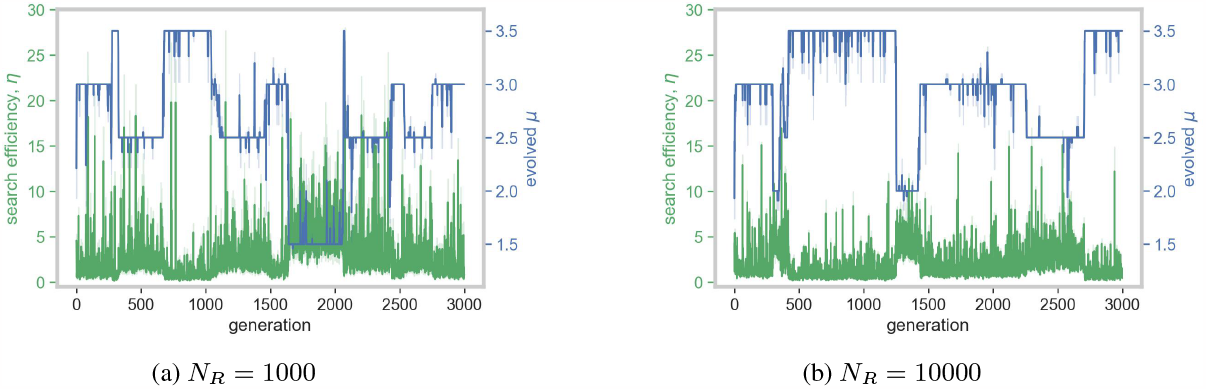
Example of an evolutionary trajectory of the mean value of a group’s *μ* (in blue) and *η* (in green) over 3000 generations for a single simulation of a group of size 10 with *α* = 10^−5^. The shaded regions indicate 68% confidence intervals of the mean estimates.

Taken together, our results suggest that explorative agents are necessary for groups to be efficient. Explorative strategies may not be beneficial for an individual if others can easily exploit their discoveries without contributing new discoveries to the group. However, these results are from search strategies where agents have fixed behavioral regimes which cause explorative agents to continue exploring for new resources even after finding a resource rather than exploiting it. We next test whether adaptive switching between exploration and exploitation can increase individual-level benefits of explorative strategies and resolve the discrepancy between what is optimal for an individual vs. the collective.

### 4.4. Adaptive switching between exploration and exploitation can balance individual and group efficiencies

In the results discussed thus far, explorative searchers find resource patches quickly but also are quick to exit the found patches before fully exploiting them. Social learning can put them at a disadvantage by allowing others to exploit their discoveries. However, if explorers could adaptively exploit their own discoveries, they would be able to increase their payoffs. To test this idea, we created agents who could use local information to adaptively decide when to continue exploiting a patch and when to exit it to explore for more patches. This decision mimicked the widely-observed *area-restricted search* (ARS) strategy. ARS, thus, enabled explorative foragers to individually exploit found resources by engaging in local search when resources are found.

We found that with ARS groups evolved to be more explorative. With ARS, explorative agents could adaptively exploit a patch before exiting it, which meant that exploitative agents could not effectively scrounge discoveries from their explorative counterparts. This effect caused the mean *μ* values to decrease below 2 (Fig. 5a) with fewer exploiters selected (Fig. 6). In addition, fluctuations in *μ* were smaller with ARS than without it (Fig. 4 (bottom row)) because explorative strategies had substantially higher payoffs than exploitative ones.

**Figure 4:**
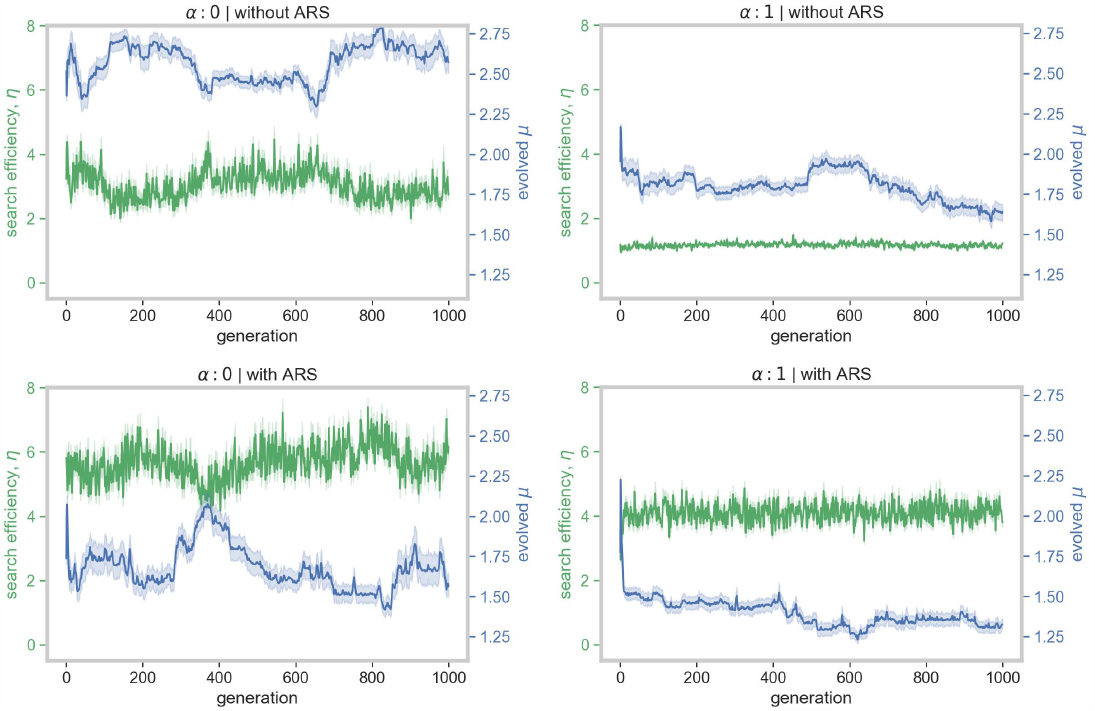
Time-average of trends over all simulations in efficiency, *η* and Lévy exponent, *μ* over 1000 generations for groups of size, 10 and *N*_*R*_ = 1000. We calculated the moving average over the window size of 20 generations to show clearer trends. The shaded regions indicate 95% confidence intervals of the mean estimates.

**Figure 5:**
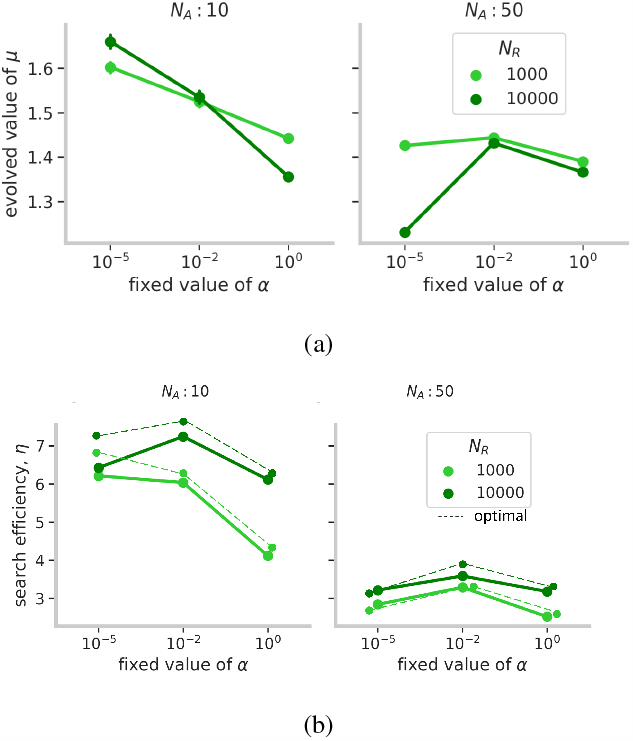
***(a)*** Mean estimates of the evolved values of Lévy exponents (*μ*) from the area-restricted search (ARS) model for different levels of resource density (*N*_*R*_), group size (*N*_*A*_) and social learning (*α*). ***(b)*** Corresponding mean estimates of the group search efficiencies (*η*) of the evolved groups. The averages were taken over the last 10 generations out of a total of 3000, for every parameter combination. Error bars indicate 95% confidence intervals.

**Figure 6:**
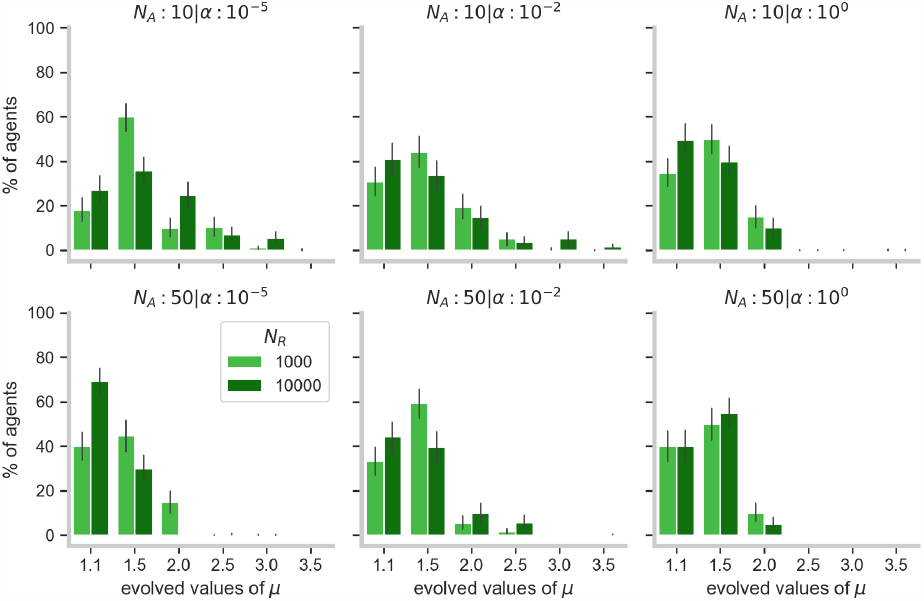
Distribution of the evolved values of Lévy exponents (*μ*) for different levels of resource density (*N*_*R*_), group size (*N*_*A*_) and social learning (*α*). These data represent group compositions over the last 20 generations out of a total of 3000. *α* values of 10^−5^, 10^−2^, 10^0^ correspond to high, intermediate, and no levels of social learning. *μ* → 1 corresponds to explorative search strategy, while *μ* → 3.5 corresponds to exploitative search. The error bars indicate 95% confidence intervals.

In terms of group-level search efficiencies, we found that the higher proportions of explorative agents allowed the groups to maximize their search efficiencies (Fig. 5b) close to the theoretical optimum (shown in dashed lines in Fig. 5b) as estimated from the model reported by Garg et al. (2022) (For details, see Supplementary Results and Fig. S7). Moreover, unlike the non-ARS condition where social learning led to a decrease in search efficiency over generations (Fig. S2), foragers using ARS became more efficient (Fig. S3).

However, the previously noted relationship between the level of exploration and social learning continued to hold. Overall levels of exploration decreased with social learning but in large groups, agents became highly explorative. This effect was strongest with dense patches, where groups were more explorative than even in the case of no social learning (*α* = 0). This effect occurs because ARS is more effective with dense patches than sparser ones and, thus, it was beneficial for agents to spread out and decrease competition [Di Bitetti and Janson, 2001].

Finally, ARS with *μ* → 1 (long steps interspersed with informed short steps) can resemble non-ARS Lévy search with *μ* ≈ 2. The difference between the two lies in whether information about the environment drives the decision to switch between exploring and exploiting or whether the switch occurs randomly. Results from invasion analyses (Fig. S6) show that, on average and across conditions, Lévy walks with *μ* ≈ 2 were least likely to be invaded by exploiters or explorers. Taken together, our results indicate that search strategies that balance exploration and exploitation, whether by informed or random decisions, can be evolutionary stable with social learning.

## 5 Discussion

Foraging in groups has many benefits and costs that can affect an individual’s search strategies [Krause et al., 2002, Ranta et al., 1993, Cvikel et al., 2015]. Optimal search models have mostly focused on solitary foraging, where a forager has to balance explorative and exploitative modes of search to discover and harvest resources. Our model of general collective foraging shows that the evolution of individual search strategies, in terms of exploration and exploitation, can depend upon social interactions among foragers in groups. The model also shows that differences in explorative tendencies in independent search can result in frequency-dependent payoffs that may not always improve group search efficiency. Instead, individual-level selection can lead to discrepancies between individual and collective search goals [Leonard and Levin, 2022] by favoring exploitative strategies that are less efficient in the long run. We found that informed search strategies (here, ARS) that use environmental information to guide search decisions can restore and stabilize the balance between exploration and exploitation at both individual and group levels. Such a balance can improve both individual and collective search efficiencies, and result in the evolution of more optimal search groups.

Previous studies have shown that random search strategies that can balance search for close and distant resources can be optimal for individual foragers [Bartumeus et al., 2014, Garg and Kello, 2021, Bénichou et al., 2011]. Our results show that this balance may not always be optimal in groups where information about found resource locations is shared between foragers. Instead, we found that the optimal strategy for a searcher depends upon what strategies others adopt: if group-level exploration is high, then it can be beneficial to be exploitative, and vice-versa. However, if groups become too exploitative, search can be inefficient for both groups and individuals. We found that the presence of social learning requires a concomitant increase in the role of cognitive processing in individual search strategies to maintain and enhance search efficiency. In other words, in the face of competition, individuals need to adaptively switch between exploration and exploitation. Our results add to previous studies showing how the addition of simple heuristics to random-walk models can greatly increase search efficiencies when compared to purely random search models [Hills et al., 2013, Ross and Winterhalder, 2018, Reynolds, 2009, Hein and McKinley, 2012]. Results also suggest that minimizing the role of cognitive processing will tend to underestimate the extent to which non-local search is explorative. Indeed, random search models might be especially insufficient to capture real-world behavior under conditions of threat and competition [Mobbs et al., 2018]. Under such conditions that pose significant opportunity costs, foragers may not be able to afford random search and may also evolve more complex strategies with complex communication, memory, and environmental sensing [Hecker and Moses, 2015].

Evidence for simple heuristics like ARS has been found in humans and other animals [Pacheco-Cobos et al., 2019, Wiesner et al., 2012, Dorfman et al., 2022]. Our results suggest that ARS may confer an evolutionary advantage to social foragers by increasing their share of resources. Furthermore, prior work suggests that ARS patterns (i.e., short, exploitative bouts alternating with longer excursions) can closely resemble Lévy patterns of *μ* ≈ 2 and be more efficient than strategies where the switch between exploration and exploitation is random. Some works have shown that Lévy patterns of *μ* ≈ 2 can be optimal in social contexts, too, over multiple aspects. They can help balance cooperation and competition [de Jager et al., 2011] and lead to superdiffusive collective motions in swarms ([Ariel et al., 2015]). Santos et al. Santos et al. [2009] also showed that Lévy patterns can help balance collective cohesiveness and dispersion, and optimize collective efficiency in groups with leaders and followers. Given that our results show that on average, *μ* ≈ 2 is the strategy that is least likely to be invaded across all conditions (Fig. S6), our results support the possibility that processes that can balance explorative bouts with exploitative efforts [Kölzsch et al., 2015, Sims et al., 2019, Campeau et al., 2022] can be bet-hedging strategies to stabilize equilibria in groups. Whether search strategies with *μ* ≈ 2 emerge from environmental factors or are a result of evolved Lévy processes, the level of exploration or exploitation that they give rise to will be subject to varying payoffs and selection. Our results highlight basic relations between individual and collective search that can be expected to occur in real biological and social search dynamics.

In our previous paper [Garg et al., 2022], we showed that collective search can be efficient if individual foragers are highly explorative and quickly find new resources while being selective in their social learning to exploit and collectively exploit clusters of resources found. Herein we showed social learning requires foragers to be more intelligently exploitative to protect their efforts from scrounging. Other studies on social foraging have shown that foragers may use certain strategies to increase their finder’s share [Di Bitetti and Janson, 2001, Vickery et al., 1991]. For instance, capuchin monkeys maintain large distances between each other while searching for food, in order to harvest a sufficient amount of food before others join in. Likewise, in our model, explorative foragers with ARS were better at finding resources and maintaining distances between each other. This strategy may also be in line with predictions from the ideal free distribution theory of animal foraging which suggests that foragers in a group distribute themselves across a resource landscape in ways that minimize competition while maximizing energetic returns [Fretwell, 1969, 1972]. Different social systems may employ other mechanisms to protect from scrounging and more generally promote exploration, such as social prestige or synchronized food-sharing in hunter-gatherers [Winterhalder, 1996]. Multi-level selection, where foragers compete with each other but also face group-level pressures to cooperate, can also give rise to competitive groups that maintain a high proportion of explorers for higher efficiencies.

Our results show that the level of social learning practiced by the group can determine the search strategies of individuals. Reliance on social learning increased the benefits of exploitative search strategies but in larger groups, greater opportunities for social learning favored explorative strategies. Similar to our model’s predictions, studies of bees have shown that the level of exploration practiced by the bees increases with group size, due to competition for limited resources [Grüter and Hayes, 2022]. However, our model assumes that the tendency or ability to make foraging decisions on based on social learning is independent of the individual search strategy. These search features may be correlated [Kurvers et al., 2010], in which case, groups would benefit from a mix of asocial explorers who find resource patches and social exploiters who harvest a found patch. The movement or space-use patterns can also affect how foragers acquire information (e.g., due to speed-accuracy trade-offs) [Spiegel and Crofoot, 2016] or how well they can communicate with each other [Roeleke et al., 2022]. Further investigations can modify the model’s assumptions to test the effects of explorative strategies on group-level efficiencies and general adaptability.

Individuals in many animal groups can consistently differ from each other in their search strategies, especially in terms of explorative behavior [Reader, 2015, Mehlhorn et al., 2015], that in turn can affect group-level behaviors related to foraging such as cohesiveness, flocking, risk-taking[Aplin et al., 2014, Ioannou and Dall, 2016, Ward et al., 2004, Dyer et al., 2009, Burns and Dyer, 2008]. Our work adds to this discussion and shows instead how the differences between individuals in their explorative tendencies may be affected by physical and social environment features. We show that the differences in the movement speed and patch discovery can lead to dynamics similar to the classic producer-scrounger models [Barnard and Sibly, 1981, Caraco and Giraldea, 1991]. Our results support previous empirical work that has shown explorative and exploitative foraging behavior to be density-dependent [Sokolowski et al., 1997, Greene et al., 2016]. Our model also suggests that these differences may not always be adaptive, and a high proportion of exploiters can decrease the mean fitness of a population. Although a mix of explorers and exploiters in a group was not theoretically optimum in our model, that may not always be the case in the natural world. In some socially foraging species, where individual fitness is tightly linked with that of the group, explorative scouts do not optimize their finder’s share and instead abandon food sources after discovering and recruiting other workers [Grueter and Leadbeater, 2014, Liang et al., 2012]. In many natural conditions, explorative strategies may have additional risks (such as predation, high search costs, or reduced attention to social cues) that could decrease efficiencies in groups composed solely of explorers. For instance, in a variable environment, if the most rewarding option is associated with high risk, then explorers that continue searching for better options would be selected against [Arbilly et al., 2011]. Furthermore, maintaining a mix of diverse search strategies may be especially helpful in variable and uncertain resource environments [Dingemanse et al., 2004].

Studies on optimal search strategies have largely focused on the individual-level and how the physical environment can shape the strategies. The present work shows that the studies on optimal search strategies need to account for the social environment, as well. We show that an individual’s search behavior is constrained by both the physical and social environment, and can, in turn, shape the group and its capabilities. Our model also highlights how the differences between random and informed search strategies can lead to important consequences on both individual- and collective-level search efficiencies, especially under competitive foraging. Future models on social foraging should account for the role social information plays in shaping individual preferences and search behavior, and how social learning is affected by independent search behavior. In this paper, we used the explorative-exploitative movements to highlight these trade-offs but social foraging models can easily be extended to other aspects of search behavior such as optimal departure time, optimal travel time that is formulated within the Marginal Value Theorem framework [Davis et al., 2022]. For instance, decisions to explore or exploit can be driven by the perceived value of resource patches that can be modeled to take into account the costs and benefits of both the physical and social context [Silston et al., 2021]. There may also be additional trade-offs posed by being in a group, such as maintaining group cohesiveness, and leader-follower dynamics, that affect these decisions [Santos et al., 2006].

The implications of our results are not limited to foraging for resources but extend to collective problem-solving and action, where independent searchers use social information to find solutions to a problem. Studies on collective problem-solving and search can benefit from investigating how individual search behavior within a group might be influenced by the strategies adopted by others. For instance, if some group members are risk-prone and explorative, then others might prefer searching for less-riskier solutions, minimizing their search costs, and prefer to improve upon solutions found by others. Or, being a part of a group may dilute risk and embolden group members to explore novel and risky solutions to a problem [Camisón-Zornoza et al., 2004]. There may be additional incentives for the exploration of novel solutions or ideas, for example, patents, social prestige, and other rewards associated with innovations [Giraldeau et al., 2017]. Our results also suggest that studies on collective behavior should consider the discrepancies between individual and collective goals, costs, and benefits [Leonard and Levin, 2022]. Further, investigating how social learning and communication evolve in tandem with individual search strategies under different contexts can shed light on general aspects of collective behavior and sociality.

## 6 Author Contributions

K.G, C.T.K. and P.E.S contributed to the conception and design of the model, wrote and edited the paper; K.G. coded and analyzed the model.

## 7 Data Availability

Code is available on this GitHub repository - https://github.com/ketikagarg/collective_foraging.

## 8 Acknowledgements

We thank the editors and reviewers for their helpful suggestions.

## 9 Supplementary Methods

### 9.1 Resource Environment Generation

We generated resource environments (Fig. S1) using a power-law distribution growth model. The space was initialized with 20 seed resources placed in random locations. Additional resources were placed such that the probability of a resource appearing a distance *d*_*r*_ from previously placed resources was given by

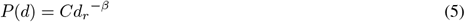

where, *d*_*min*_ ≤ *d*_*r*_ ≤ *L*, ≤ *d*_*min*_ = 10^−3^ is the minimum distance that an agent could move and *L* = 1 is the normalized size of the grid. *C* is a normalization constant required to keep the total probability distribution equal to unity, such that

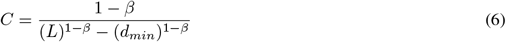

*β* determines the spatial distribution of resources other than the resource seeds, such that *β* → 1 resembled a uniform distribution and *β* → 3 generated an environment where resources were tightly clustered. In this paper, we set *β* = 3 to generate distinct resource patches.

### 9.2 Evolutionary Invasion Analysis

We systematically tested whether a population of composed of agents with a given *μ* can be invaded by a mutant agent with another value of *μ*. To perform invasion analysis, we simulated homogeneous populations of a *μ*_*resident*_ ranging between 1 and 3.5, and for each of the resident populations, we added a mutant with another value of *μ*_*mutant*_ between 1 and 3.5. Each of these populations was run for only one generation (i.e., they did not evolve), and we tested the likelihood of a given *μ*_*mutant*_ invading *μ*_*resident*_ by calculating an invasion index, *i*, for a pair of two different values of *μ* where:

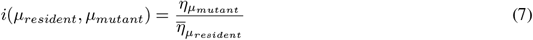

## 10 Supplementary Results

### 10.1 Area-restricted search

In order to compare the results from the evolutionary model with the Garg 2022 model, we implemented the ARS condition in the 2022 model, too.

We found that the addition of an area-restricted search component to the model changed the optimal search efficiencies both at the individual and the group levels. With pure Lévy walks and no social learning, *μ* = 2 was optimal search behavior because search movements are a mix between long, explorative steps and short, exploitative ones. But with ARS, explorative agents can adaptively switch to exploitative steps after encountering resources and may not need short steps otherwise. Indeed, with ARS added, *μ* = 1.1 resulted in randomly oriented explorative movements mixed with informed exploitative bouts and optimal search efficiencies (Fig. S7). The mix also increased search efficiencies (Fig. S7) for all values of *μ* and *α* when compared to pure Lévy walks, and selective social learning was more beneficial with ARS than without it.

**Fig. S1:**
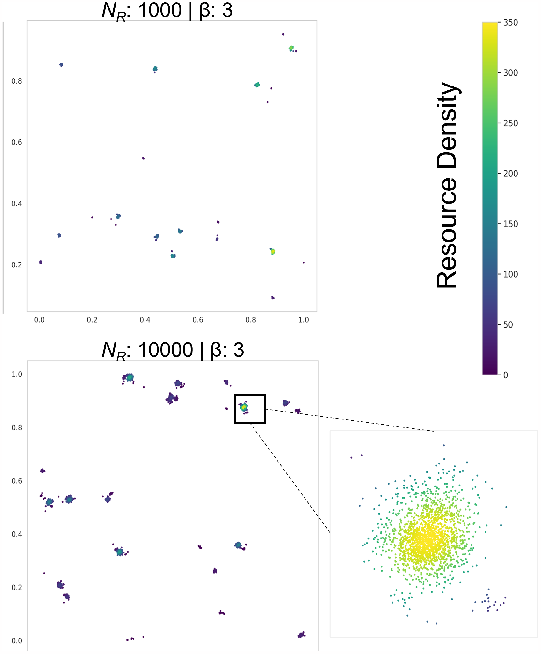
Examples of the resource distributions generated by the power-law growth algorithm with *β* = 3. The color-map indicates the density estimates (calculated using Gaussian Kernel Density Estimation) of resources present at a location.

**Fig. S2:**
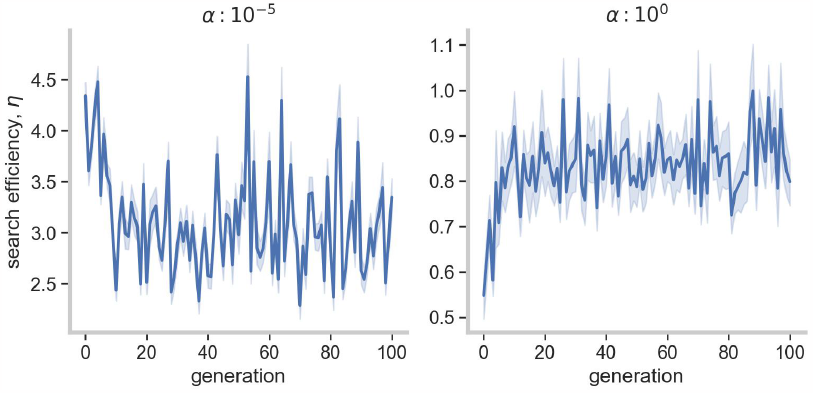
Average trends in collective search efficiency over initial generations, for the non-ARS condition, group size of 10 and resource density of 10,000. The figure shows that in the absence of social learning (*α* = 10^0^), search efficiency of the group increases, but with social learning (*α* = 10^−5^), search efficiency decreases. Note the different y-axes limits for the two panels.

**Fig. S3:**
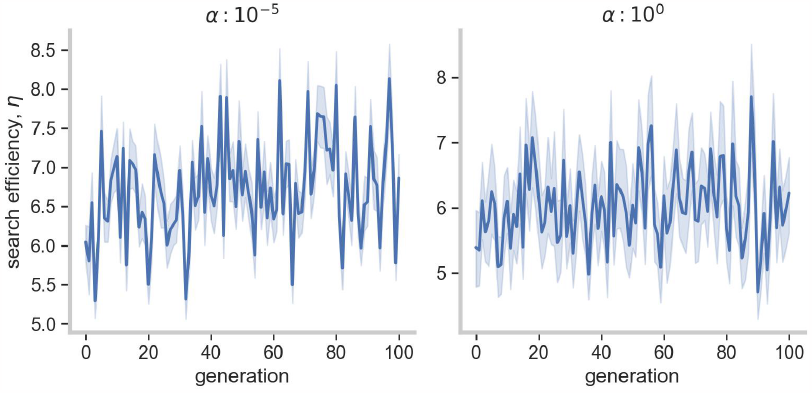
Average trends in collective search efficiency over initial generations, for the ARS condition, group size of 10 and resource density of 10,000. The figure shows that with and without social learning, search efficiency of the group increases over time. However, with social learning (*α* = 10^−5^), the increase is more substantial than without (*α* = 10^0^). Note the different y-axes limits for the two panels.

**Fig. S4:**
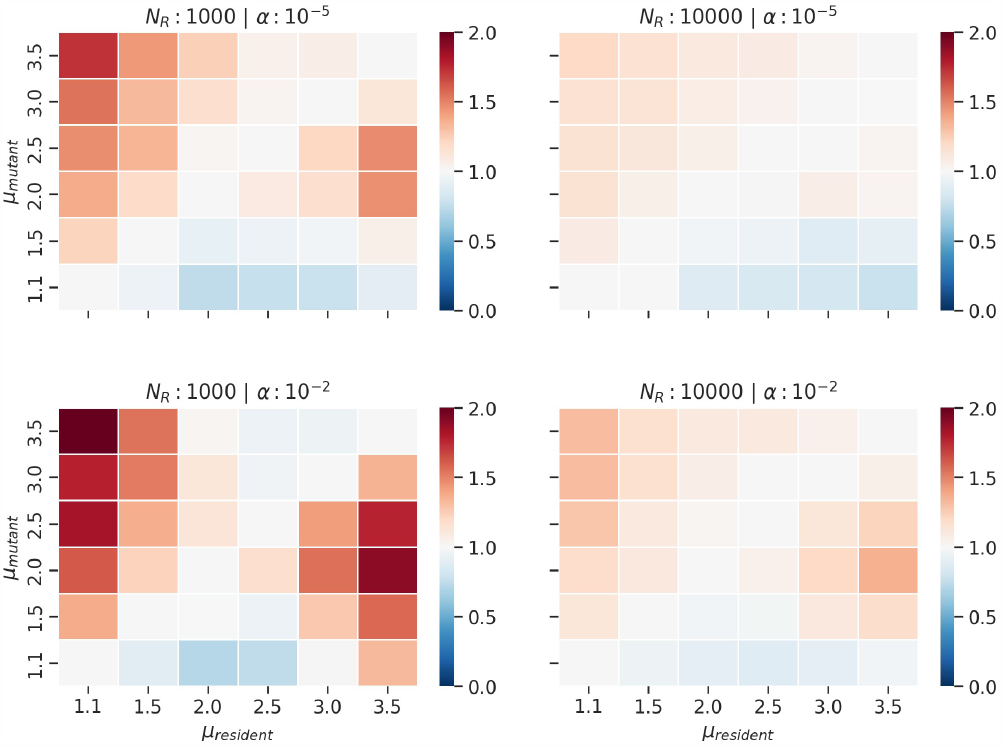
Mean estimates of invasion index for groups of size 10, different levels of resource density (*N*_*R*_) and social learning (*α*) over 500 simulations. Index values greater than 1 imply that the mutant *μ* will be over to invade the resident *μ*.

**Fig. S5:**
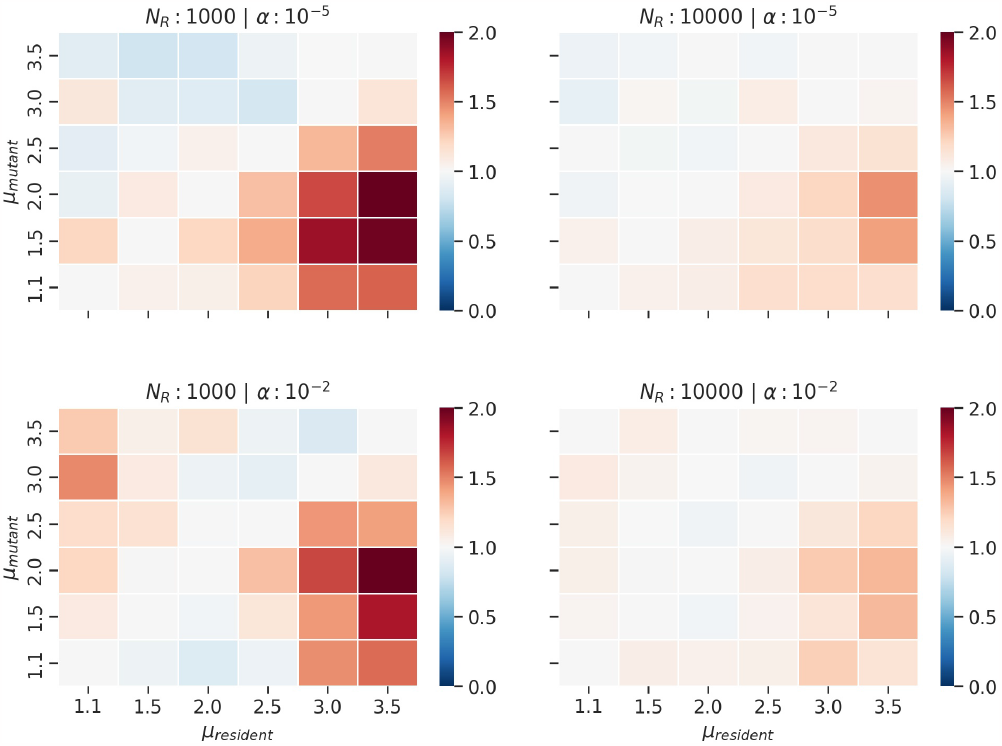
Mean estimates of invasion index for groups of size 50, different levels of resource density (*N*_*R*_) and social learning (*α*) over 500 simulations. Index values greater than 1 imply that the mutant *μ* will be over to invade the resident *μ* and are represented by red hues.

**Fig. S6:**
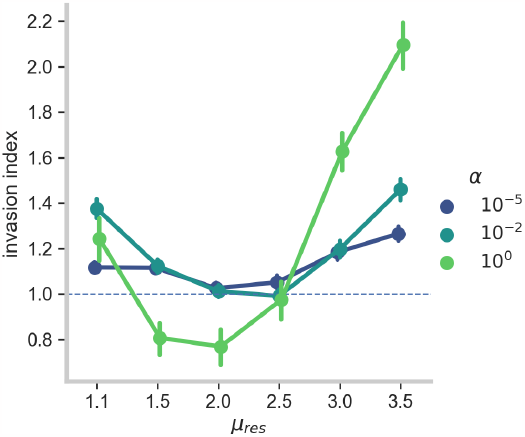
Overall mean estimates of invasion index for all groups of size 10 and 50, different levels of resource density (*N*_*R*_) and different *μ*_*mut*_. Different colors indicate the three levels of social learning (*α*). This plot shows that on an average among all the different conditions, *μ*_*res*_ ≈ 2 is least likely to be invaded by other strategies.

**Fig. S7:**
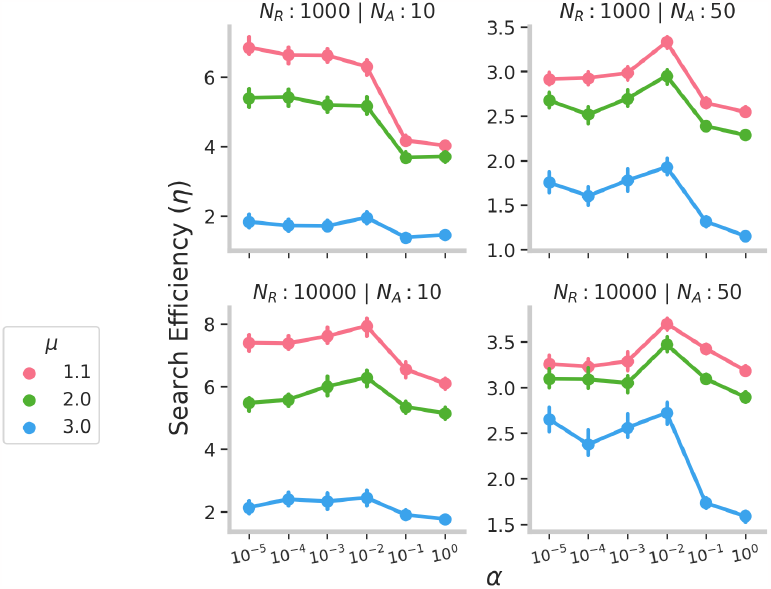
Group search efficiency *η* for the ARS model as a function of social selectivity parameter *α*, Lévy exponent *μ*, resource density *N*_*R*_. Error bars indicate 95% confidence intervals. Note the different Y-axes limits.

